# Conformational changes in Arp2/3 complex induced by ATP, WASp-VCA and actin filaments

**DOI:** 10.1101/164319

**Authors:** Sofia Espinoza-Sanchez, Lauren Ann Metskas, Steven Z. Chou, Elizabeth Rhoades, Thomas D. Pollard

## Abstract

We used fluorescence spectroscopy and electron microscopy to determine how binding of ATP, nucleation-promoting factors (NPF), actin monomers and actin filaments change the conformation of Arp2/3 complex during the process that nucleates an actin filament branch. We mutated subunits of *Schizosaccharomyces pombe* Arp2/3 complex for labeling with fluorescent dyes at either the C-termini of Arp2 and Arp3 or ArpC1 and ArpC3. We measured Förster resonance energy transfer (FRET) efficiency (ET_eff_) between the dyes in the presence of the various ligands. We also computed class averages from electron micrographs of negatively stained specimens. ATP binding made small conformational changes of the nucleotide binding clefts of the Arp subunits. WASp-VCA, WASp-CA, and WASp-actin-VCA changed the ET_eff_ between the dyes on the Arp2 and Arp3 subunits much more than between dyes on ArpC1 and ArpC3. Ensemble FRET detected a different structural change that involves bringing ArpC1 and ArpC3 closer together when Arp2/3 complex bound actin filaments. Each of the ligands that activates Arp2/3 complex changes the structure in different ways, each leading progressively to fully activated Arp2/3 complex on the side of a filament.

## Introduction

Forces generated by the actin cytoskeleton are crucial to numerous cellular processes including cell migration and endocytosis (Pollard and Borisy 2003, Goley and Welch 2006). However, spontaneous nucleation of actin filaments is kinetically unfavorable due to the rate-limiting formation of actin dimers and trimers (Cooper, Buhle et al. 1983, Pollard, Blanchoin et al. 2001, Sept and McCammon 2001). Cells use proteins to overcome the kinetic barrier to nucleation, but must regulate these proteins carefully (Pollard, Blanchoin et al. 2001, Chhabra and Higgs 2007).

The first actin nucleating protein discovered was Arp2/3 complex (Machesky, Atkinson et al. 1994), a complex of actin-related proteins Arp2 and Arp3 and five other supporting subunits: ARPC1 (p40), ARPC2 (p34), ARPC3 (p21), ARPC4 (p20), and ARPC5 (p16). Arp2/3complex is intrinsically inactive, because the supporting subunits hold Arp2 and Arp3 too far apart to nucleate a daughter filament (Robinson, Turbedsky et al. 2001, Goley and Welch 2006). When active Arp2/3 complex binds to the side of a “mother” actin filament, which serves as a base for initiating a new “daughter” filament as a branch at a characteristic 70° angle (Mullins, Heuser et al. 1998). Activation is thought to depend on a large conformational change that brings Arp2 and Arp3 about 30 Å closer together (Robinson, Turbedsky et al. 2001, Goley and Welch 2006) arranged like two subunits of an actin filament(Aguda, Burtnick et al. 2005 Egile, Rouiller et al. 2005 Rouiller, Xu et al. 2008, Xu, Rouiller et al. 2012) as observed in 3D reconstructions of electron micrographs of branch junctions (Rouiller, Xu et al. 2008).

*In vitro*, three factors cooperate to stimulate the nucleation activity of the complex: actin monomers, a mother filament and a nucleation-promoting factor (NPF) (Marchand, Kaiser et al. 2001, Dayel and Mullins 2004, Zhang, Wu et al. 2005). However, the contributions of each factor to the conformational changes required to activate Arp2/3 complex remain unclear.

Members of the Wiskott-Aldrich syndrome protein (WASp) family of proteins are NPFs (Machesky, Mullins et al. 1999, Rohatgi, Ma et al. 1999, Winter, Lechler et al. 1999, Yarar, To et al. 1999). These proteins share a conserved C-terminal VCA motif with one or two verprolin homology (V) sequences, a central (C) sequence and an acidic (A) sequence. The V motif binds monomeric actin, the C motif interacts with both actin and Arp2/3 complex in an independent but mutually exclusive manner, and the A motif binds exclusively to Arp2/3 complex (Marchand, Kaiser et al. 2001). Free VCA is mostly unfolded/extended (Marchand, Kaiser et al. 2001) but assumes secondary structure when bound to Arp2/3 complex (Panchal, Kaiser et al. 2003) or actin monomers (Chereau, Kerff et al. 2005, Pollard 2007). Importantly, Rho-family GTPases regulate NPFs, thus linking upstream cellular signals to actin polymerization (Ma, Rohatgi et al. 1998, Mullins and Pollard 1999, Rohatgi, Ma et al. 1999, Bompard and Caron 2004).

Several studies aimed to characterize how NPFs and mother filaments promote the conformational change and activate Arp2/3 complex. Goley *et al*. (Goley, Rodenbusch et al. 2004) investigated the effect of different ligands on Arp2/3 complex structure by ensemble F′rster Resonance Energy Transfer (FRET) using fluorescent proteins. They fused YFP to the C-terminus of ArpC3 and CFP to the C-terminus of ArpC1. ATP, GST-WASp-VCA and Wasp-CA each increased energy transfer modestly. Actin filaments did not change the transfer efficiency (unpublished data in Goley *et al*.). Compelling evidence implicates VCA in conformational changes (Martin, Xu et al. 2005 Rodal, Sokolova et al. 2005 Xu, Rouiller et al. 2012, Hetrick, Han et al. 2013), but a pathway specifying the contributions of each ligand to the final conformation is still missing.

Several lines of evidence provide indirect information about the conformational changes that activate Arp2/3 complex, but neither the structural changes nor their relation to ligand binding are firmly established. Inspection of the crystal structure of inactive Arp2/3 complex suggested that a block of the structure including Arp2, ARPC1, ARPC5and part of ARPC4 rotates about 30° with respect to the rest of the complex, bringing Arp2 and Arp3 together (Figure 1 Lower panel) (Robinson, Turbedsky et al. 2001). This model was supported by steered molecular dynamics simulations (Dalhaimer and Pollard 2010), 3D reconstruction of electron micrographs of single particles of Arp2/3 complex with NPFs (Xu, Rouiller et al. 2012) and a combination of EM and computational analysis (Egile, Rouiller et al. 2005). Others proposed that Arp2 moves independently with respect to the other subunits of the complex (Irobi, Aguda et al. 2004) (Aguda, Burtnick et al. 2005) (Figure 1 Middle panel). This model was supported by statistics-based docking of the crystal structure of the inactive Arp2/3 complex into a 3D reconstruction of the branch junction from electron tomograms (Rouiller, Xu et al. 2008) and refinement by molecular dynamics (MD) simulations (Pfaendtner, Volkmann et al. 2012).

**Figure 1.**
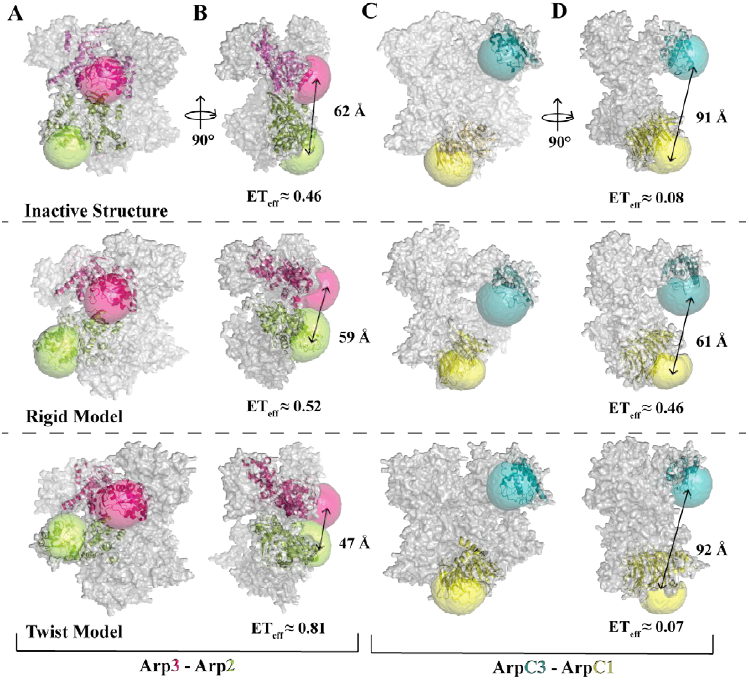
Models of three different conformations of Arp2/3 complex with accessible volume simulations of the fluorophores (colors) and distances between labeling sites. *Top row:* Crystal structures of inactive Arp2/3 complex (Coordinates from pdb files 1K8K and 4JD2)). *Middle row:* Model of active Arp2/3 complex based on reconstruction from electron micrographs of actin filament branch junctions, fitting the crystal structure into the density and refinement by all atom molecular dynamics simulations (Pfaendtner et al, 2012). *Bottom row:* Model of partially activated Arp2/3 complex from steered molecular dynamics simulations (Dalhaimer and Pollard, 2010). (**Columns A, B**) Front and side views of the three models of Arp2/3 complex with Accessible Volume (AV) simulations (FPS software *(Kalinin, Peulen et al. 2012)*) of fluorophores on the C-termini of subunits Arp3 (*magenta*) and Arp2 (*green*). (**Columns C, D**) Front and side views of the three models of Arp2/3 complex with AV simulations of fluorophores on the C-termini of subunits ArpC1 (*yellow*) and ArpC3 (*cyan*). Inter-dye distances (bold) were calculated using the AV mean fluorophore positions. ET_eff_ values were calculated using Equation 2 (See materials and methods).

Here we present single molecule and bulk FRET measurements to characterize how binding of ATP, WASp-VCA, WASp-CA, and WASp-VCA with bound monomeric actin influence the conformation of Arp2/3 complex. We used pairs of small fluorescent dyes conjugated to cysteine residues at the C-termini of Arp2, Arp3, ARPC1 and ARPC3. ATP binding has only minimal effects on single molecule FRET and electron microscopy of single particles, showing that ATP binding closes the nucleotide-binding cleft of Arp2. Single molecule and bulk FRET measurements are consistent with NPF binding favoring a partial rotation Arp2 with associated subunits to bring it closer to Arp3. Ensemble FRET measurements show that binding to an actin filament not only causes this rotation but also rearrangement of other subunits.

## Results

### Design of smFRET sensors for Arp2/3 complex conformations

To investigate the effects of different activating factors on the conformation of Arp2/3 complex, we modified the genome of fission yeast *Schizosaccharomyces pombe* to add cysteine residues to the C-termini of subunits of Arp2/3 complex for labeling with fluorescent dyes (Figure 1). We started with a strain modified with the mutation ARPC4 C167S that removes the only reactive cysteine of Arp2/3 complex (Beltzner and Pollard 2008). Strain Arp2_cys_-Arp3_cys_ has single cysteines at the C-termini of Arp2 and Arp3, and strain ArpC3_cys_-ArpC1_cys_ has single cysteines at the C-termini of ArpC1 and ArpC3. Both strains are viable. After purification on three columns, Arp2/3 complex from both strains was >99% pure (Figures 2A and Supplemental Figure 4) Purified complexes were labeled with Alexa-Fluor-488-C_5_-maleimide (Alexa 488) and/or Alexa-Fluor-594-C_5_-maleimide (Alexa 594), using a 2-fold molar excess of the donor Alexa 488 and a 10-fold molar excess of the acceptor Alexa 594. R0 for this FRET pair is 54 Å (Schuler, Lipman et al. 2002). On average each complex had 0.10 donor dye (Alexa 488) and 1.8 acceptor dyes (Alexa 594) (Figure 2A) distributed among three subpopulations: donor-only, acceptor-only, and double-labeled.

**Figure 2.**
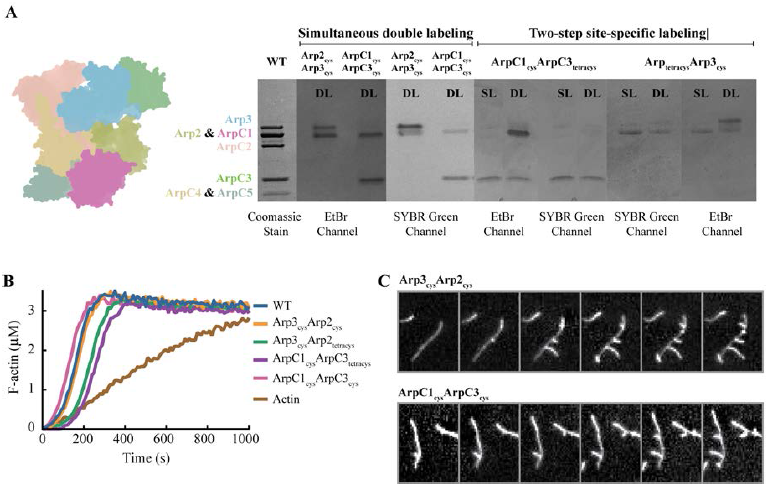
Labeled Arp2/3 complexes nucleate and branch actin filaments. (**A**) Gel electrophoresis of purified and labeled Arp2/3 constructs with data from seven different conditions aligned. Constructs ArpC1_cys_-ArpC3_cys_ and Arp3_cys_-Arp3_cys_ were labeled simultaneously with the FRET pair Alexa 488 and Alexa 594. Constructs ArpC1cys-ArpC3tetracys and Arp3_cys_-Arp2_tetracys_ were labeled sequentially with the FRET pair FlAsH-EDT2 followed by Alexa 568. *Left* Color-coded representation of Arp2/3 complex crystal structure. First gel lane: gel stained with Coomassie blue with representative purified Arp2/3 complex. Other lanes show the fluorescence of each complex with UV transillumination at 312 nm. For visualizing Alexa 488 and FlAsH setting for SYBR GREEN imaging were selected in the EDAS 290 Kodak Molecular Imagin Software and for visualizing Alexa 568 and Alexa 594 the Ethidium Bromide Channel was selected. DL (Double Labeled), SL (Single Labeled). (**B**) Time course of polymerization of 3 μM actin monomers (10% pyrene-labeled) with 100 nM unlabeled or labeled Arp2/3 complex and 500 nM WASP VCA at room temperature in KMEI buffer (50 mM KCl, 1 mM MgCl_2_, 1 mM EGTA, 0.1 mM ATP, 1 mM DTT, 10 mM imidazole, pH 7.0). (**C**) TIRF microscopy at intervals of ^*^ s of 1 μM actin monomers (20% labeled with Alexa-488), 500 nM WASP VCA, and 100 nM of the indicated Arp2/3 complex polymerized at room temperature in TIRF buffer (50 mM KCl, 1 mM MgCl_2_, 1 mM EGTA, 0.1 mM ATP, 1 mM DTT, 10 mMimidazole (pH 7.0), 0.02 mM, 15mM glucose, 0.02 mg/mL catalase, and 0.1 mg/mL glucose oxidase with 0.25% methylcellulose).

The labeling sites were chosen to give strong FRET signals and to distinguish between the two main hypothetical mechanisms by which Arp2/3 complex might achieve an active conformation. Accessible Volume (AV) calculations (Kalinin, Peulen et al. 2012) gave the range of motion of fluorophores conjugated to the C-termini of the four Arp2/3 complex subunits in crystal structures (Figure 1) and estimates of inter-dye distances in inactive Arp2/3 complex: 60Å between the dyes on the C-termini of Arp2 and Arp3 with a predicted FRET efficiency (ET_eff_) of 0.46; and 90 Å between dyes on the C-termini of ArpC1 and ArpC3 with a predicted ET_eff_of 0.08 (Figure 1). Partial activation by the twist model moves Arp2 and Arp3 15Å closer together with only a minimal change in the separation of the C-termini of ArpC1 and ArpC3, which are near the axis of rotation. On the other hand, activation by the *rigid model* would move the C-termini of ArpC1 and ArpC3 closer by 30 Åwithout significantly changing the separation of the C-termini of Arp2 and Arp3.

Bulk polymerization assays with pyrenyl-actin showed that in the presence of 500 nM VCA, both labeled Arp2/3 complexes promoted actin polymerization in a concentration dependent manner undistinguishable from wild type Arp2/3 complex (Figure 2B). Total internal reflection fluorescence (TIRF) microscopy showed that both labeled complexes formed actin branches with VCA as well as wild type Arp2/3 complex (Figure 2C). Thus neither the addition of C-terminal cysteines nor labeling with Alexa dyes impaired the activity of Arp2/3 complex.

### Energy transfer efficiencies between labels on Arp2/3 complex by single-molecule FRET

We measured FRET of individual double-labeled Arp2/3 complexes at a concentration of ~75pM. We fit primary smFRET histograms with Gaussian distributions and calculated the mean position and width of the non-zero peaks (Table S3).

Arp2/3 complex labeled on Arp2 and Arp3 had two ET_eff_ peaks (Figure 3). We tested the identity of the ET_eff_ ≈ 0 peak using Alternative Laser Excitation (ALEX) to determine if any ET_eff_ peaks were buried in this area (Figure 3B). ALEX showed that the first peak comes exclusively from donor-only labeled complexes due to their lack of responsiveness from a 561 nm laser (Kapanidis, Laurence et al. 2005) (Figure 3B, C). The main peak had a mean ET_eff_ of 0.48 ± 0.02 (n: 9) and a width of 0.14 ± 0.03 (n: 9). This agrees with the predicted ET_eff_ of 0.46 for this pair according to the crystal structure. Under ideal conditions, these ET_eff_ values correspond to a distance of 60 ± 10 ÅThe Supplemental Information has details and fluorophore controls.

**Figure 3.**
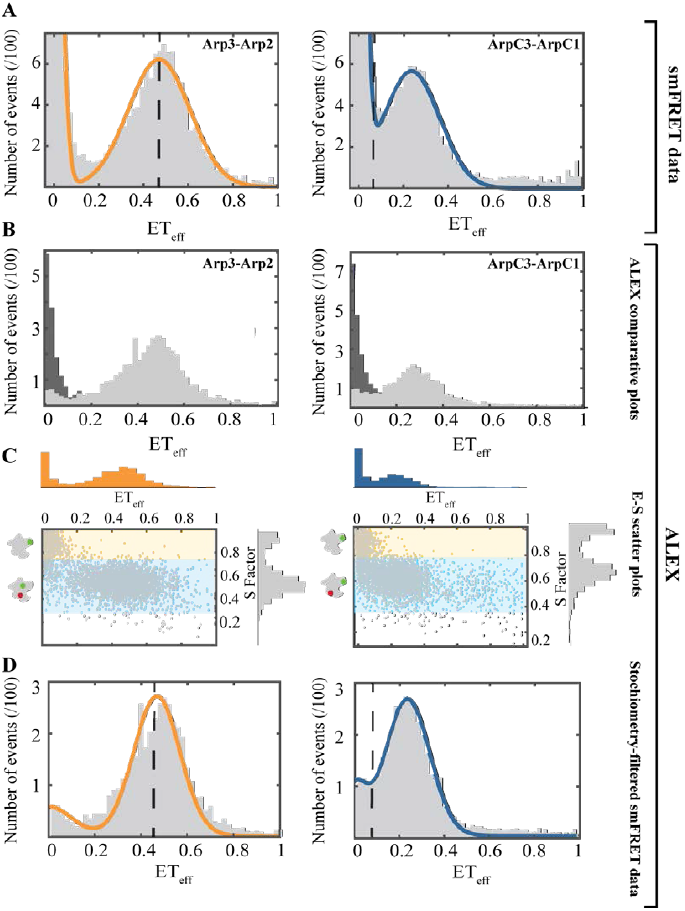
Single-molecule FRET and Alternating-Laser Excitation (ALEX) measurements show single populations of labeled Arp2/3 complex. *Left panels*: Arp2_cys_-Arp3_cys_ construct. *Right panels:* ArpC1_cys_-ArpC3_cys_ construct. Conditions: 75 pM Arp2/3 complex and 40 nM unlabeled Arp2/3 complex in KMET buffer (50 mM KCl, 1 mM MgCl2, 1 mM EGTA, 0.1 mM ATP, 1 mM DTT, 10 mM Tris-HCl, pH 7.0). (**A**) Distribution of diffusion-based single molecule FRET events with continuous wave, monochromatic excitation. (**B**) Distributions of donor-only and donor-acceptor molecules sorted by <u>A</u>lternating-<u>L</u>aser <u>Ex</u>citation (ALEX) *(Kapanidis, Laurence et al. 2005)* with donor-only events in dark grey and donor-acceptor events in light grey. (**C**) Scatter plots of ET_eff_ vs. stoichiometry (E-S) with histograms for E along the x-axis, and for S along the y-axis. Ratio E sorts species according to FRET, and ratio S sorts species according to donor-acceptor stoichiometry. Donor-only species have low E and high S. (**D**) Stoichiometry-filtered smFRET histograms using ALEX analysis to exclude donor-only species. *Orange* Fitted curve for Arp2_cys_-Arp3_cys_ construct. *Blue* Fitted curve for ArpC1_cys_-ArpC3_cys_ construct. *Dashed black lines:* Locations of ET_eff_ peaks predicted from Accessible Volume (AV) simulations based on the crystal structure of inactive Arp2/3 complex (Fig. 1 top row).

The labeled ArpC3_cys_-ArpC1_cys_ construct also had two smFRET peaks (Figure 3A), the zero peak and a peak centered at a mean ET_eff_ of 0.25AV calculations from the crystal structure predicted an ET_eff_ peak buried in the zero peak area; however, ALEX confirmed that the low efficiency peak contained only donor-only labeled complex (Figure 3B). The main peak had a mean ET_eff_ of 0.25± 0.01 (n: 7) and a mean width of 0.11 ± 0.01 (n: 7). This mean ET_eff_ is higher than the value of 0.08 predicted for this construct (Figure 1). This difference corresponds to a ~20 Åshorter distance than the predicted one from the crystal structure. The observed ET_eff_ value corresponds to an approximate distance of around 70 ± 10 Å (See Supplemental Information for details and fluorophore controls.)

### Effect of ATP on the conformation of Arp2/3 complex

A single-molecule FRET experiment showed that saturating concentrations of ATP (2 mM) did not change the smFRET histograms of either the ArpC3_cys_-ArpC1_cys_ pair (Figure 4A) or the Arp2_cys_-Arp3_cys_ FRET pair (Figure 4B). Neither the mean position nor the width of the ET_eff_ peaks changed for either pair of labeling sites.

We confirmed that ATP does not cause large conformational changes by analyzing single Arp2/3 complex molecules in electron micrographs of negatively stained specimens (Figure 4 C, D). We classified and averaged >10,000 molecules in the presence and absence of ATP. The samples were quite uniform, because the front side of Arp2/3 complex is relatively flat and most particles landed front side down on the grid.

The class averages clearly resolved Arp2 (including the nucleotide binding cleft), Arp3 (including the nucleotide binding cleft), ArpC1, ArpC3 and ArpC5 (Figure 4C). ArpC2 and ArpC4 appeared as a continuous density as expected in a projection image. In the favored view Arp3 is viewed from the top, looking down the nucleotide-binding cleft, while the front side of Arp2 is visible. The highest density is in the region where ArpC1, ArpC4 and subdomains 3 and 4 of Arp2 superimpose in the projection, consistent with crystal structures.

**Figure 4.**
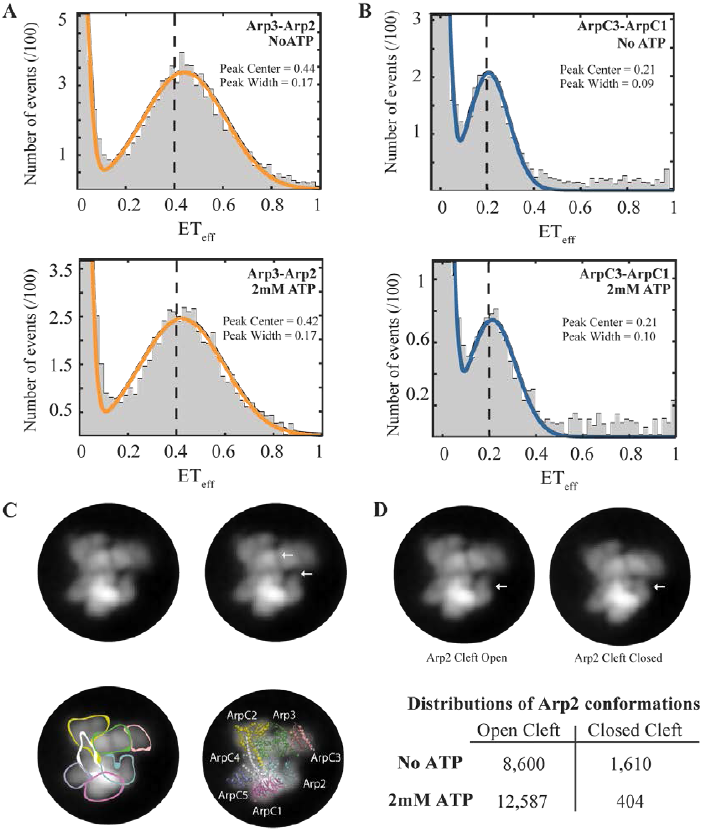
Effect of ATP on the conformation of Arp2/3 complex. (**A and B**) Single molecule FRET experiments. Conditions: 75 pM Arp2/3 complex and 40 nM unlabeled Arp2/3 complex in KMET buffer (50 mM KCl, 1 mM MgCl_2_, 1 mM EGTA, 1 mM DTT, 10 mM Tris-HCl, pH 7.0) without ATP (upper panels) or with 2 mM ATP (lower panels). (**A**) Arp2_cys_-Arp3_cys_ construct (upper panel) without ATP and (lower panel) with 2 mM ATP. (**B**) ArpC1_cys_-ArpC3_cys_ construct (upper panel) without ATP and (lower panel) with 2 mM ATP. (**C and D**) Electron microscopy in the absence and present of ATP. Samples were prepared by negative staining and imaged by transmission electron microscopy. Virtually all of the particles had the same orientation on the support film. After correction for drift, particles were classified and class averages computed.All four images are the dominant class average of the projection structure of Arp2/3 complex without ATP computed from 2066 particles. Top left, image alone. Top right, white arrows indicate the nucleotide-binding clefts of Arp2 and Arp3. Bottom left, the subunits are outlined. Bottom right, a ribbon diagram of the crystal structure of inactive Arp2/3 complex is superimposed. (D) Comparison of the dominant class averages (left) without ATP with the nucleotide-binding cleft of Arp2 open and (right) with ATP with the nucleotide-binding cleft of Arp2 closed. The projection structure of Arp2/3 complex with ATP was computed from 2705 particles. White arrows indicate Arp2 subunits. The table has the distributions of Arp2/3 complex particles classified with the nucleotide-binding cleft of Arp2 open or closed in the presence or absence of ATP.

Two class averages predominated in the samples without ATP: one with the nucleotide-binding cleft of Arp2 slightly open (84% of particles); and one with the cleft closed (16% of particles) (Figure 4D). Thus the lack of order of subdomains 1 and 2 in most crystal structures (Nolen and Pollard 2008) arises from a range of hinge angles between the two halves of Arp2. We expect the same to be true for Arp3, but the viewing angle is not favorable for detecting differences.

The class average structures in the presence of ATP were similar to the apo structures except that the Arp2 nucleotide binding cleft was closed in 97% of the particles. The distance between ArpC1 and ArpC3 was the same in the particles with open and closed Arp2 clefts.

### VCA/CA ligands increase the FRET efficiency between labels on Arp2 and Arp3 more than between ARPC1 and ARPC3

We used single-molecule FRET to study interactions of Arp2/3 complex with CA, VCA and VCA crosslinked to an actin monomer (actin-VCA). We used fluorescence correlation spectroscopy (FCS) to confirm that VCA binds Arp2/3 complex under our experimental conditions (Supplemental Figure 1). Alone in solution 20 nM labeled VCA has a τ of 0.40 ms. In the presence of 500 nM Arp2/3 complex a two-component diffusion model fit the data for labeled VCA: 58% of the population had τ _1_ = 0.91 ms like Arp2/3 complex alone in solution and the rest had τ _2_ = 0.32 ms close to the 0.40 ms τ value of free VCA. This extent of binding agrees with previously reported K_d_’s for VCA and Arp2/3 complex (Ti, Jurgenson et al. 2011).

Fluorescence anisotropy confirmed that Alexa 488-labeled VCA and CA constructs bind to Arp2/3 complex (Supplemental Figure 2). Anisotropy values were 0.11 for 50 nM VCA and 0.10 for 50 nM CA. In the presence of 3 μM Arp2/3 complex, the anisotropy values increased to 0.21 for VCA and to 0.22 for CA. These values are similar to the value 0.22 for 100 nM Alexa 488 single-labeled Arp2/3 complex. These results and previously reported K_d_s of less than 2 μM (Ti, Jurgenson et al. 2011), validated the concentrations used to saturate the complex.

Single molecule FRET experiments showed that 20 μM VCA, 20 μM CA, and 2 μM actin-VCA had different effects on the ArpC3_cys_-ArpC1_cys_ and Arp2_cys_-Arp3_cys_ constructs (Figure5). All three ligands increased the mean ET_eff_ peak value of the Arp2_cys_-Arp3_cys_ construct indicating a decrease in the inter-dye distance. VCA increased the ET_eff_ by 0.13, CA by 0.06 and actin-VCA by 0.12. All of the smFRET histograms showed two peaks, but ALEX confirmed that the ET_eff_ ≈ zero peaks come solely from donor-only labeled complexes (Figure 5 insets). In contrast, none of these ligands changed the ET_eff_ for the ArpC3_cys_-ArpC1_cys_ construct substantially. VCA and CA each increased the ET_eff_ by 0.03 and actin-VCA increased ET_eff_ by 0.02.

**Figure 5.**
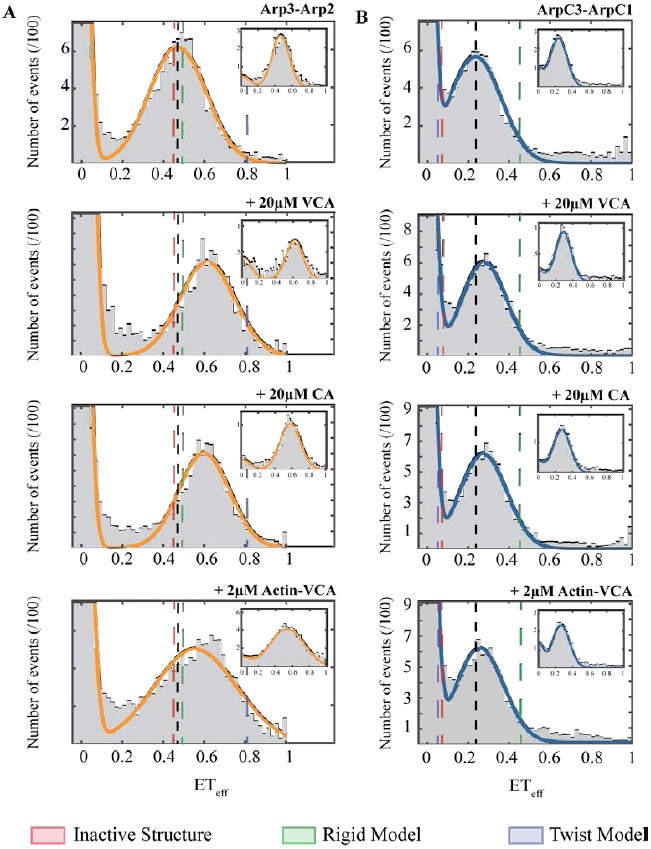
VCA/CA ligands increase the FRET efficiency between labels on Arp2 and Arp3 more than between labels on ARPC1 and ARPC3. Distributions of diffusion-based single molecule FRET events for (**A**) Arp2_cys_-Arp3_cys_ construct, and (**B**) ArpC1_cys_-ArpC3_cys_ construct. Conditions: 75 pM labeled Arp2/3 complex in KMET buffer (50 mM KCl, 1 mM MgCl_2_, 1 mM EGTA, 0.1 mM ATP, 1 mM DTT, 10 mM Tris-HCl, pH 7.0) with *top row*, 40 nM unlabeled Arp2/3 complex; *second row,* 20 μM WASP-VCA from fission yeast Wsp1p; *third row,* 20 μM WASP-CA from fission yeast Wsp1p; and *fourth row,* 2 μM actin-VCA from fission yeast Wsp1p. *Insets:* Representative stoichiometry-filtered ALEX histograms. *Gray:* Representative histograms of smFRET events collected over 60 min. *Orange:* Fitted curves for Arp2_cys_-Arp3_cys_ construct. *Blue:* Fitted curves for ArpC1_cys_-ArpC3_cys_construct. *Black* dashed line: Reference line indicating the peak center for Arp2/3 complex alone. Dashed lines: Predicted ET_eff_ peaks from *red,* the inactive crystal structure, *green*, the rigid EM model, and *blue*, the MD twist model (Fig. 1 top row).

### Effect of actin filaments on the conformation of Arp2/3 complex

Single molecule FRET experiments in solution were not possible with filamentous actin, even when the lengths of the filaments were restricted with capping protein (data not shown), so we used ensemble FRET of orthogonally double-labeled Arp2/3 complex to evaluate the effect of actin filaments on the conformation of Arp2/3 complex (Figure 6). We purified Arp2/3 complex from strains Arp3_cy_s-Arp2_tetracys_ and ArpC1_cys_-ArpC3_tetracys_. Strain Arp3_cys_-Arp2tetra_cys_ has a single cysteine at the C-terminus of Arp3 and a peptide with four cysteine residues at the C-terminus of Arp2, and strain ArpC1_cys_-ArpC3_tetracys_ has a single cysteine at the C-terminus of ArpC1 and the tetracysteine peptide at the C-terminus of ArpC3. Purified complexes (Figure 2A) were labeled using a two-step process. First, we used a 10-fold molar excess of the donor FlAsH-EDT2 and then a 7-fold molar excess of the acceptor Alexa-Fluor-568-C_5_-maleimide (Alexa 568). R0 is ~75Åfor this FRET pair (Granier, Kim et al. 2007). On average, labeling was 50% donor dye (FlAsH-EDT_2_) and 100% acceptor dye (Alexa 568) (Figure 2B, see materials and methods for further details). The C-terminal tetracysteine peptide and labeling with FlAsH dye reduced the activity of Arp2/3 complex a small amount (Figure 2B).

**Figure 6.**
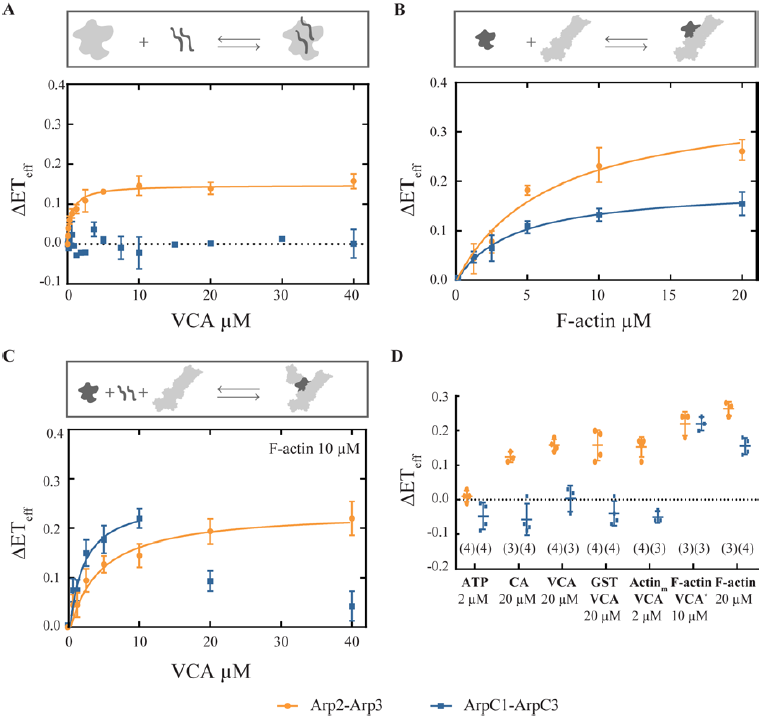
Actin filaments increase the FRET efficiency between dyes on both Arp2/Arp3 and ArpC1/ArpC3. Ensemble ET_eff_ measurements of 20 nM Arp2/3 complex labeled (*Blue)* with FlAsH on ArpC3 and Alexa 568 on ArpC1 or (*Orange*) with FlAsH on Arp2 and Alexa 568 on Arp3. The measurements in each experiment are normalized to conditions without the varied ligand. Conditions: 50 mM KCl, 1 mM MgCl_2_, 1 mM EGTA, 0.1 mM ATP, 1 mM DTT, 10 mM imidazole, pH 7.0, unless indicated otherwise. (**A**) Titration of Arp2/3 complex with Wsp1p-VCA. (**B**) Titration of Arp2/3 complex with polymerized actin with overnight incubation. (**C**) Titration of Arp2/3 complex and 10 μM polymerized actin with Wsp1p-VCA. (**D**) Interleaved scatter plot of normalized mean 㥂ET_eff_ values (± 1 SD) of labeled Arp2/3 complex in the presence of 2 mM ATP and various ligands: 20 μM Wsp1p-CA, 80 μM Wsp1p-VCA, 20 μM polymerized actin, 2 µM actinm-VCA (actin monomer convalently linked to VCA), 20 μM GST-VCA, or 10 μM polymerized actin with 10 μM VCA. Numbers of repetitions are in parentheses.

Saturating concentrations (2 mM) of ATP did not change the overall ET_eff_ of either construct (Supplemental Table 4), and the two constructs responded differently to 20 μM CA, 20 μM VCA, 20 μM GST-VCA and 2 μM actin-VCA (Figure 6 and Supplemental Table 4). All of these ligands increased ET_eff_ of Arp2_tetracys_-Arp3_cys_, but had no effect or decreased the ET_eff_ for the ArpC1_cys_-ArpC3_tetracys_ construct. These results confirm our smFRET measurements. Titration of Arp2_tetracys_-Arp3_cys_ with VCA increased ET_eff_ consistent with a Kd of 0.64 μM while the ET_eff_ of the ArpC1_cys_-ArpC3_tetracys_ construct did not change (Fig. 5A).

Twenty micromolar actin filaments increased the ET_eff_ of both ArpC1_cys_-ArpC3_tetracys_ and Arp3_cys_-Arp2_tetracys_, (Figure 6B). The actin filament concentration dependence gave K_d_s of 6.6 μM for Arp3_cys_-Arp2_tetracys_ and 4.13 μM for ArpC1_cys_-ArpC3_tetracys_.

The double-labeled ArpC1_cys_-ArpC3_tetracys_ and Arp2_tetracys_-Arp3_cys_ constructs responded differently to titration with VCA in the presence of 10 μM actin filaments (Figure 6C). The ET_eff_ of Arp2tetra_cys_-Arp3_cys_ increased up to 0.22 ± 0.03 (n = 3) at 40 μM VCA. A hyperbola fit to this data gave a K_d_ of 4.43 μM. On the other hand, the ET_eff_ of ArpC1_cys_-ArpC3_tetracys_ increased only up to 10 μM VCA (K_d_ of 2.03 μM) and declined at higher concentrations of VCA.

## Discussion

The crystal structure of inactive Arp2/3 complex suggested that a large conformational change is required to reposition the Arps to initiate the daughter filament (Robinson, Turbedsky et al.2001), so many methods have been used to investigate how nucleotides, nucleation promoting factors and mother filaments promote the active conformation. This discussion explains how our spectroscopic measurements and electron microscopy provide new insights about these conformational changes. Our work extends the conclusions from previous experiments using X-ray crystallography, F′rster energy transfer, chemical crosslinking, radiation footprinting and mass spectrometry, electron microscopy of single particles and branch junctions, and molecular dynamics simulations.

### Inactive Arp2/3 complex has one major conformation

High-resolution crystal structures (extending to 2.0 Å) provide the most detailed information about the structure of inactive Arp2/3 complex. Crystal packing forces can influence protein conformations, but three different crystal forms of bovine Arp2/3 complex (Robinson, Turbedsky et al. 2001, Ti, Jurgenson et al. 2011, Luan and Nolen 2013) and crystals of fission yeast Arp2/3 complex (Nolen and Pollard 2008) do not differ substantially. This conformation was stable during 10 ns of atomistic, molecular dynamics simulations of inactive Arp2/3 complex except for small fluctuations in local regions, such as the nucleotide binding clefts of the Arp subunits (Pfaendtner and Voth 2008, Dalhaimer and Pollard 2010).

The class averages from our electron micrographs of negatively stained fission yeast apo-Arp2/3 complex confirm the presence of one major conformation with a small range of different hinge angles between the two halves of Arp2. In contrast to the uniformity of these structures, an early EM study reported the coexistence of three different conformations of negatively stained budding yeast and bovine Arp2/3 complexes (Rodal, okolova et al. 2005). All three of their conformations are similar to our class averages for apo-Arp2/3 complex, except that Arp2 seems to be missing or disordered in their open conformation. More details are visible in our class averages, because we analyzed more particles and used movies to correct for drift.

Our single molecule FRET data along two dimensions of the complex with small dyes on the C-termini of Arp2 and Arp3 or ArpC1 and ArpC3 supports the consensus that inactive fission yeast Arp2/3 complex in solution at room temperature largely has one conformation. Energy transfer between dyes on the C-termini of Arp2 and Arp3 match the value predicted from the crystal structure (Figure 3A). The ET_eff_ between ArpC1 and ArpC3 is higher than expected, so the labeled residues appear to be 20 Åcloser together than expected from the crystal structure. However, the expected distance is greater than the limit for accurate measurements, and the dyes may have preferred conformations not captured by the AV calculations or the anisotropy/lifetime measurements.

Both constructs have single peaks in the single molecule ET_eff_ histograms, suggesting that Arp2/3 complex adopts a preferred conformation in solution on the millisecond timescale of our measurements. This is consistent with predictions from MD simulations, which indicated that the intrinsic fluctuations of the complex are substantially more rapid than our sampling timescale, which would result in a single ET_eff_ value due to the averaging of the signal over time (Dalhaimer and Pollard 2010). The peak widths were generally consistent with shot noise estimates and other potential non-conformational contributions, with the notable exception of the Arp2_cys_-Arp3_cys_ bound to actin-VCA. This construct has a peak that is substantially broader than the upper boundary expected for shot noise (Supplemental Table S3), which may suggest some conformational heterogeneity or slower-timescale dynamics compared to other conditions.

### ATP binding results in local conformational changes

Crystal structures (Nolen, Littlefield et al. 2004) (Nolen and Pollard 2007), all-atom MD simulations (Dalhaimer, Pollard et al. 2008) and radiation footprinting with mass spectrometry (Kiselar, Mahaffy et al. 2007) all showed that central cleft of Arp3 closes around bound ATP or ADP, although ATP bound to subdomains 3 and 4 of Arp2 did not stabilize the disordered subdomains 1 and 2 in the crystal structures. Bound nucleotide did not change other parts of Arp2/3 complex in these studies. Three-dimensional reconstructions from electron micrographs of frozen-hydrated and negatively stained Arp2/3 complex from budding yeast and *Acanthamoeba* were homogeneous and indistinguishable from the crystal structures except for the presence of subdomains 1 and 2 of Arp2 (Xu, Rouiller et al. 2012). Our class averages of electron micrographs of fission yeast Arp2/3 complex show that ATP stabilizes subdomains 1 and 2 and closes the cleft of Arp2.

In agreement with this evidence, ATP binding to Arp2/3 complex produces little or no change in FRET efficiency between dye probes on Arp2 and Arp3 or on ArpC1 and ArpC3 in our single molecule experiments. On the other hand, a pioneering study of recombinant human Arp2/3 complex using ensemble FRET measurements of bulk samples with YFP fused to the C-terminus of ArpC1 and CFP fused to the C-terminus of ArpC3 concluded that “nucleotide binding promotes a substantial conformational change in (Arp2/3) complex” (Goley, Rodenbusch et al. 2004). They reported FRET/CFP ratios, which were determined by dividing the acceptor peak emission by the donor peak emission. The FRET/CFP ratio changed by 0.11 upon binding to ATP. We recalculated ET_eff_ using equation 2 (See Materials and Methods) and obtained an ET_eff_ increase of only ~0.02, which is consistent with our data. Although more complicated to prepare, our Arp2/3 complex labeled with small dye molecules was fully active, while Arp2/3 complex fused to fluorescent proteins was 20 times less active than wild type Arp2/3 complex.

ATP binding in the nucleotide cleft of Arp3 can release the C-terminus of Arp3 from the barbed end groove (Rodnick-Smith, Liu et al. 2016). This rearrangement might change the energy transfer efficiency of a dye on the C-terminus. However, we did not observe a change, so the average position of the dye does not change significantly after the release of the C-terminus of Arp3. Thus, binding of ATP may trigger local conformational changes that do not cause FRET changes between our specific probes.

### Binding of nucleation promoting factors reorients Arp2 and Arp3

Several studies aimed to characterize how NPFs and mother filaments promote the conformational change that activates Arp2/3 complex. We agree with the general assumption that full activation brings Arp2 and Arp3 into contact like successive subunits along the short-pitch helix of an actin filament, as observed in 3D reconstruction of the branch junction (Rouiller, Xu et al. 2008). Our single-molecule FRET data and observations by five other methods agree that nucleation promoting factors favor a conformation with the Arps moved part of the way toward the fully active conformation, which has only been observed in the branch junction.

Steered molecular simulations of Arp2/3 complex involved applying a spring-like force between Arp2 in the inactive crystal structure and its position in the branch junction determined by electron microscopy (Dalhaimer and Pollard 2010). During atomistic molecular dynamics simulations, the force moved Arp2 towards the active target position, along with several associated subunits. In 12 separate experiments a block of protein consisting of Arp2, ARPC1, ARPCand part of ARPC4 rotated together by twisting a pair of alpha-helices that connect ArpC4 to ArpC2. In 11 experiments steric interference stalled this rotation after about ~30°. In one simulation the block of protein with Arp2 moved past this barrier further toward Arp3 in an actin filament-like structure. These results suggest a pathway for the initial part of the conformational change. Reaching the position of Arp2 in the branch junction would require a further rotation of 15° around a different axis. Assuming that NPF binding is responsible for the first partial movement, actin filament binding might favor the final conformational change. Five different approaches provide further information about now nucleation-promoting factors influence the conformation of Arp2/3 complex.

Goley *et al*. (Goley, Rodenbusch et al. 2004) used FRET between YFP fused to the C-terminus of ArpC3 and CFP fused to the C-terminus of ArpC1 to investigate the effect of NPFs on Arp2/3 complex. Both GST-WASp-VCA and WASp-CA changed the FRET/CFP ratio by 0.23. Our recalculation using equation 2 gives an ET_eff_ increase ~0.05 upon binding to WASp-CA. This small change agrees with our measurements of ET_eff_ between dyes on the C-termini of ArpC1 and ArpC3 in single molecule FRET experiments. Our ensemble experiments did not detect this change. Thus the distance between the C-termini of ArpC1 and ArpC3 changes little when CA or VCA binds, as expected from their locations close to the axis of rotation in steered MD simulations.

Binding of VCA or CA increased ET_eff_ between probes on the C-termini of Arp2 and Arp3 without changing the width of these distributions, so the population average switches to a conformation with the two Arps closer together. The rotation observed in the steered MD simulations can explain this change. On the other hand, if the FRET change were due to the release of the C-terminus of Arp3, the dye would sample more positions, which would likely broaden the distribution of FRET efficiencies, which was observed with actin-VCA binding Arp2/3 complex. Thus Arp2/3 complex with bound actin-VCA may sample a wider range of conformations than with bound CA or VCA. Two electron microscopy studies also showed that nucleation promotion factors favor a “closed” conformation of *Acanthamoeba*, bovine and budding yeast Arp2/3 complex with the Arps closer together than inactive Arp2/3 complex (Rodal, Sokolova et al. 2005 Xu, Rouiller et al. 2012).

Chemical crosslinking of Arp2 and Arp3 provides the most direct evidence for VCA favoring a conformation with the Arps close together. Rodnick-Smith *et al*. (Rodnick-Smith, Liu et al. 2016) substituted single cysteines into budding yeast Arp2 and Arp3 that are adjacent when the Arps are in the short-pitch conformation but not inactive Arp2/3 complex. The bifunctional, 8 Å crosslinker bis-maleimidoethane covalently links the cysteine-substituted Arp2 and Arp3 in minutes and even faster with N-WASP-VCA. Thus the budding yeast Arp2/3 complex must spontaneously visit the short-pitch conformation, as reflected by its ability the nucleate daughter filaments without nucleation promoting factors. The crosslinked complex nucleates actin filaments without nucleation promoting factors, so the crosslink stabilizes an activated state of Arp2/3 complex.

Radiation footprinting and mass spectrometry showed that WASp VCA binding to bovine Arp2/3 complex triggers an allosteric conformational change involving only subunits Arp2 and Arp3. The authors proposed the domain rearrangement of Arp2 and Arp3 could result in a closed conformational state (Kiselar, Mahaffy et al. 2007). This contrasts with what happens upon ATP binding, as this change is not localized within a specific region of the subunits (Kiselar, Mahaffy et al. 2007).

### Effect of actin filament binding on the conformation of Arp2/3 complex

Binding a pair of nucleation promoting factors (Padrick, Doolittle et al. 2011, Ti, Jurgenson et al. 2011, Boczkowska, Rebowski et al. 2014) and the side of an actin filament (Machesky, Mullins et al. 1999) fully activates Arp2/3 complex to nucleate a daughter filament. Two dimensional (Egile, Rouiller et al. 2005) and three dimensional (Rouiller, Xu et al. 2008) reconstructions of branch junctions established that Arp3 and Arp2 are arranged along a short-pitch actin filament helix and are the first two subunits of the daughter filament. Fitting the crystal structure of Arp2/3 complex into the density between the mother and daughter filaments required movement of Arp2 and adjacent subunits. Pfaendtner et al. (Pfaendtner, Volkmann et al. 2012) used the electron microscopy reconstruction and the crystal structure of Arp2/3 complex as the starting point for large-scale MD simulations of the branch junction including 31 proteins (~3 million atoms) for 175 ns. These simulations included 13 subunits of mother filament, 11 subunits of daughter filament and the 7 subunits of Arp2/3 complex. The relatively low RMSD of ~4.3 Å of final Arp2/3 complex structure supported the overall positions of the subunits in the initial EM model. Moreover, the simulation showed that an extensive network of salt bridges between both elements reinforces binding of Arp2/3 complex to the mother filament.

Independently Goley et al. used molecular dynamics simulations and protein-protein docking simulations (Goley, Rammohan et al. 2010) to model interactions between the ARPC2 and ARPC4 heterodimer and the side of an 8 subunit actin filament. The model obtained by this approach was very similar (RMSD of5.9 Å) to the models based on electron microscopy.

Our ensemble FRET measurements with dye probes on Arp2 and Arp3 or ArpC1 and ArpC3 showed that the ET_eff_ of both constructs increases when they bind actin filaments. Thus, the relative positions ArpC1 and ArpC3 change in addition to the change between Arp2 and Arp3 induced by VCA. Goley et al. (Goley, Rodenbusch et al. 2004) did not detect a change in the transfer efficiency between fluorescent proteins fused to ArpC1 and ArpC3 (data not published), so the fluorescent proteins might have interfered with important interactions between the mother filament and ARPC1 or ARPC3 subunits (Pfaendtner, Volkmann et al. 2012) or they may not have allowed the slow reaction to proceed to completion.

### Proposed conformational changes

Overall our results suggest that both a rotation of Arp2 and associated subunits and some further rearrangement of subunits contribute to activation of Arp2/3 complex as proposed by Xu *et al*. (Xu, Rouiller et al. 2012) and Egile *et al*. (Egile, Rouiller et al. 2005) to explain reconstructions of branch junctions from electron micrographs. Egile *et al*. proposed a combined 30 Å rotation of Arp2 and Arp3 and a slight adjustment of the overall complex (≈5 Å) to fit Arp2/3 complex into EM images of actin branches (Egile, Rouiller et al. 2005).

Arp2/3 complex is inactive in solution. According to our measurements in solution, Arp2/3 complex has predominantly a single, inactive conformation that depends on interactions with VCA and actin filaments to nucleate a daughter filament. ATP binding promotes closure of the nucleotide clefts of both Arps (Nolen, Littlefield et al. 2004, Kiselar, Mahaffy et al. 2007, Dalhaimer, Pollard et al. 2008, Pfaendtner and Voth 2008) and destabilizes the C-terminus of Arp3 (Rodnick-Smith, Liu et al. 2016), but these changes do not activate the complex

VCA binding favors a conformation with Arp2 closer to Arp3 but with the distance between ArpC1 and ArpC3 unchanged. A 30° rotation of the block of structure including Arp2 (Dalhaimer and Pollard 2010) can explain these changes, the structures observed by electron microscopy (Fig. 7) and chemical crosslinking observed between Arp2 and Arp3 (Rodnick-Smith, Liu et al. 2016). The free energy from VCA binding is used to reach this activated state, but the mechanism is not yet clear. Bound VCA increases the affinity of Arp2/3 complex for the mother filament 20-fold (Ti, Jurgenson et al. 2011), presumably because the conformation of activated complex better matches the side of the filament.

**Figure 7.**
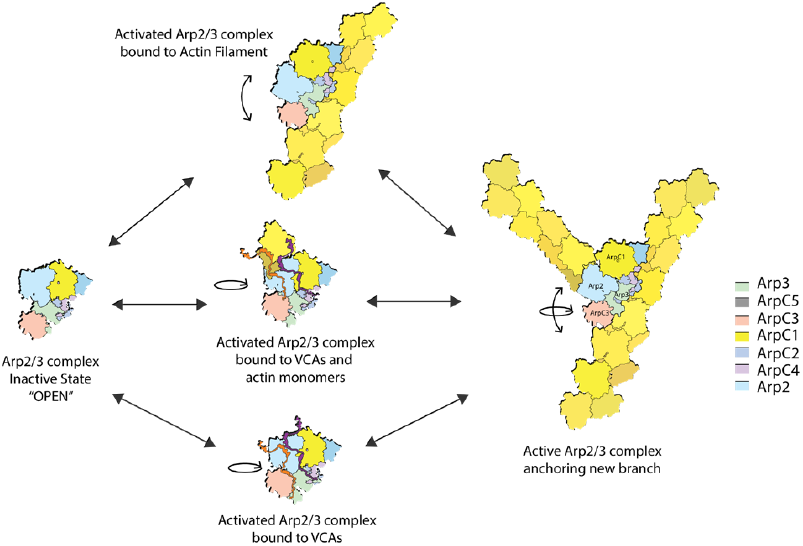
Model of conformational changes associated with Arp2/3 complex activation. The conformation of inactive Arp2/3 complex is an open state with separated Arp2 and Arp3. Binding of VCA or actin-VCA promotes a rotational movement to an activated intermediate conformation that favors binding to the side of a mother filament. Upon binding to a mother filament, Arp2/3 complex experiences a further conformational change. These two movements can occur independently or in combination to fully activate Arp2/3 complex.

Binding to mother filaments buries 9100 Å2 of Arp2/3 complex and likely provides the free energy change required to reach the fully active state that nucleates the assembly of the daughter filament. Given the FRET change we observed between dyes on ArpC1 and ArpC3, interaction with the mother filament is likely to reposition subunits in addition to the Arps. More structural studies are needed to provide higher resolution structures of the intermediates and the branch junction.

## Materials and methods

### Cloning and mutation of S. pombe Arp2/3 complex

We added a single reactive cysteine to the C-termini of Arp2, Arp3, ArpC1 or ARPC3 or a peptide with four cysteines (CCCC) on the C-termini of ArpC3 or Arp2 by transforming a DNA fragment encoding the extra residues fused to a nourseothricin (NAT) selection marker into the G418-resistant C167S strain of *S. pombe* (Bahler, Wu et al. 1998). The C167S *S. pombe* strain has the only reactive cysteine in native Arp2/3 complex replaced by serine (Beltzner and Pollard 2008). For double mutants, we crossed two different single mutant strains on SPA5 plates and separated the spores by tetrad dissection. After tetrad dissection, we grew the spores on YE5S agar plates at 25°C for 1 day and then replica-plated colonies onto YE5S-G418/NAT plates. PCR amplification and DNA sequencing confirmed strains with the desired mutations. Single and double mutants were compared with WT cells for growth at 23, 30 and 32°C. Supplemental Table S5 gives the sequences of the primers used for these steps.

### Protein Purification and Labeling

*S. pombe* Arp2/3 complex: We purified Arp2/3 complex from fission yeast by three chromatography steps: GST-VCA affinity column (GE Healthcare Life Sciences), ion exchange on a MonoQ 55 0 column (GE Healthcare Life Sciences), and size exclusion on a HiLoad Superdex 200 16/600 column (GE Healthcare Life Sciences) (Ti et al, 2011). We measured the concentration of the purified protein by absorption at 290 nm (ε = 139 030 M^-1^ cm^-1^). Purity was verified by SDS-PAGE and staining with Coomassie blue (Figures 2A and S4). After gel filtration, ArpC1cys-ArpC3cys and Arp2cys-Arp3cys constructs were exchanged by washing four times with 15 ml aliquots of low salt buffer (50 mM KCl, 10 mM imidazole, 1 mM MgCl_2_, 1 mM EGTA, 0.1 mM ATP, pH 7) using a 30K Amicon Ultra centrifugal filter unit. We labeled mutant Arp2/3 complexes with Alexa-Fluor-488-C_5_-maleimide (Alexa488) or/and Alexa-Fluor-594-C_5_-maleimide (Alexa594) (Thermofisher) in low salt buffer. For single mutants we used 10x molar excess of dye and for double mutants, 10x fold molar excess of acceptor (Alexa594) and 2x fold molar excess of donor (Alexa488). Incubating the protein for 2 min with 5x fold molar excess of 5-dithio-bis-(2-nitrobenzoic acid (DTNB) before labeling with the dyes reduced unspecific labeling. Immediately after this pretreatment, the protein was transferred to a second microcentrifuge tube which contained the dye(s) and enough volume of high salt buffer (2 M KCl, 10 mM imidazole, 1 mM MgCl_2_, 1 mM EGTA, 0.1 mM ATP, pH 7) to bring the salt concentration up to 170 mM KCl. Proteins were incubated 45 min at 4°C and buffer was exchanged into low salt buffer + 1 mM DTT using a 30K Amicon Ultra centrifugal filter unit. After gel filtration, ArpC1cys-ArpC3tetracys and Arp3cys-Arp2tetracys constructs were buffer exchanged into FlAsH labeling buffer (170 mM KCl, 10 mM imidazole, 1 mM MgCl_2_, 1 mM EGTA, 0.3 mM ATP, 0.1 mM BAL, pH 7) using a 30K Amicon Ultra centrifugal filter unit. We labeled both mutant Arp2/3 complexes overnight at 4°C with 10x molar excess 4,5-Bis(1,3,2-dithiarsolan-2-yl)-3,6-dihydroxyspiro[isobenzofuran-1(3H),9-[9H]xanthen]-3-one (FlAsH-EDT2) (Santa Cruz Biotechnology). Protein complexes were buffer exchanged into low salt buffer using a 30K Amicon Ultra centrifugal filter unit. Before labeling with 10x molar excess of Alexa 68 dye, complexes were incubated for 2 min in 5x fold molar excess of DTNB. Immediately after, the protein was transferred to a second microcentrifuge tube containing the dye and enough high salt buffer to raise the salt concentration to 170 mM KCl. Under these conditions, incubation with DTNB prevented Alexa 568 labeling the tetra cysteine peptide.

Figure 2A shows that Alexa 594 does not label the tetra cysteine tag. Proteins were incubated 45 min at 4°C and then exchanged into low salt buffer with 1 mM DTT using a 30K Amicon Ultra centrifugal filter unit. Remaining free dye was removed from all the four constructs using a 40K ZebaTM spin desalting column (Thermofisher) equilibrated with low salt buffer with 1 mM DTT. Labeling efficiency was determined by absorption at λ 49 nm (ε 73,000 M^-1^ cm^-1^) for Alexa 488, λ594 nm (ε 92,000 M^-1^ cm^-1^) for Alexa594, λ 538 nm (; 70,000 M^-1^ cm^-1^) for FlAsH-EDT2 (Granier, Kim et al. 2007), and λ 578 nm (ε 91,300 M^-1^ cm^-1^) for Alexa586. Absorption values were corrected for dye absorption at λ 280 nm, λ 400 nm, and λ 495 nm or λ 594 nm when applicable. We used freshly made complexes for spectroscopic measurements.

VCA and CA constructs: We expressed *S. pombe* Wsp1p-VCA (497Q–574D), Wsp1p-VCAcys (496C–574D), and Wsp1-CA (541SC–574D) as GST fusions in plasmid pGV67 in Rosetta (DE3) pLysS *E. coli* cells (Novagen). We purified the proteins by DEAE anion exchange chromatography and glutathione Sepharose 4B affinity chromatography. Following cleavage of VCA/CA from GST by TEV protease, VCA and CA were separated from GST by ion exchange chromatography on a RESOURCE Q column (GE Healthcare Life Sciences) and size exclusion chromatography on a HiLoad Superdex 75 16/600 column (GE Healthcare Life Sciences). We measured concentrations of unlabeled VCA and CA by absorption at 280 nm (ε = 5690 M^-1^ cm^-1^) (Ti, Jurgenson et al. 2011) and verified purify by SDS-PAGE and staining with Coomassie blue (Figure S4). We labeled a cysteine added to the N terminus of VCA and CA using a 10x molar excess of Alexa-Fluor-488 C5-maleimide overnight at °C in labeling buffer (20 mM Tris-Cl, 100 mM NaCl, 1 mM EDTA, pH 8.0). Labeled constructs were separated from free dye by size exclusion on a HiLoad Superdex 75 16/600 column equilibrated with gel filtration buffer (20 mM Tris-Cl, 100 mM NaCl, 1 mM EDTA, 1 mM DTT, pH 8.0). Labeled constructs were used in fluorescence anisotropy and FCS experiments. Labeling efficiency was determined by absorption at 495 nm (ε 73 000 M^-1^ cm^-1^). Absorption values were corrected for dye absorption at λ 280 nm and λ 400 nm.

Actin: We purified actin from chicken breast muscle acetone powder (MacLean-Fletcher and Pollard 1980) by one cycle of polymerization and depolymerization followed by size exclusion chromatography on Sephacryl S-300 in G-buffer (2 mM Tris–HCl, 0.2 mM ATP, 0.1 mM CaCl_2_ and 0.5 mM DTT, pH 8.0). The concentration of actin was measured by absorption at 290 nm (ε 26 600 M^-1^ cm^-1^). Monomeric actin was polymerized in labeling buffer (50 mM PIPES, 50 mM KCl, 0.2 mM ATP, 0.2 mM CaCl_2_, pH 7) and labeled on Cys374 with 7x molar excess of pyrene-iodoacetamide (Pollard 1984) or 10x molar excess Alexa-Fluor-488-C5-maleimeide (Mahaffy and Pollard 2006) overnight at 4°C. Labeled actin was depolymerized using G-buffer, clarified and monomers were purified by size exclusion chromatography on Sephacryl S-300 in G-buffer. Purity was verified by Coomassie blue staining SDS-PAGE (Figure S4). Labeling efficiency was determined by absorption at λ 344 nm (ε 22,000 M^-1^ cm^-1^) for pyrene-iodoacetamide and at λ 495 nm (ε 73,000 M^-1^ cm^-1^) for Alexa 488. Absorption values were corrected for dye absorption at λ 280 nm and λ 400 nm.

### Accessible Volume (AV) calculation

We calculated Accessible Volumes (AV) for the conjugated dyes using the FPS software (Kalinin, Peulen et al. 2012). Parameters used to generate AV clouds were length 20, width 4.5, and dye radius 3.5 Å for the donor fluorophore (Alexa 488) (Kalinin, Peulen et al. 2012) and length 20, width 4.5, and dye radius 4 Å for the acceptor fluorophore (Alexa 594) (Metskas and Rhoades 205). Inter-dye distances in these models were calculated using the AV mean fluorophore positions.

### Crosslinking Actin to VCA

cys-VCA was dialyzed overnight against 2 L labeling buffer (2 mM Tris-Cl, 0.1 mM CaCl_2_, 1 mM NaN_3_, 0.1 mM ATP, 2 mM TCEP-HCl, pH 8). cys-VCA (25 μM) was incubated with 18.2 mM N,N′-m-phenylenedimaleimide (PDM) for 4 h at 4°C. Precipitated PDM was pelleted by centrifuging for 10 min at 2,5 00 g. The supernatant was dialyzed three times against 1 L of labeling buffer for 1 h at 4°C. Meanwhile, actin monomers were dialyzed against 1 L of labeling buffer for 1 h at 4°C and then 2 L for 2 h. A 10x fold molar excess of PDM-cys-VCA was incubated withactin overnight at 4°C. Crosslinked products were purified by a three-step column purification: gel filtration on a HiLoad Superdex 75 16/60 column, ion exchange on a MonoQ 10/100 column, and a second round of size exclusion on a HiLoad Superdex 75 16/60 column (Ti, Jurgenson et al. 2011). Purity was verified by SDS-PAGE and staining with Coomassie blue (Figure S4).

### Actin Polymerization Assays

We tested the functional integrity of our modified Arp2/3 complexes using pyrene actin polymerization assays (Cooper, Walker et al. 1983). Reactions contained 3.5 μM actin monomers (10% pyrene-labeled), 50 nM Arp2/3 complex, 500 nM WASP–VCA in KMEI buffer (10 mM imidazole, 50 mM KCl, 1 mM EGTA, 1 mM MgCl_2_, 1 mM DTT). Polymerization was monitored over time by fluorescence with excitation at λ 365 nm and emission at λ 407 nm using Spectramax Gemini XPS Microplate Spectrofluorometer.

### Ensemble FRET measurements

Measurements were made in a PTI Alpha-scan fluorimeter (Photon Technology International) (Marchand, Kaiser et al. 2001). FlAsH-EDT_2_ single-labeled (Arp2_FlAsH_ and ArpC3_FlAsH_) and FlAsH-EDT_2_ and Alexa 568 double-labeled complexes (Arp2_FlAsH_-Arp3_Alexa568_ and ArpC3_FlAsH_-ArpC1_Alexa568_) were diluted to fixed labeled concentrations of 20 nM in KMEI buffer. All experiments had 0.2 mM ATP present, except when testing the effect ATP on constructs where ATP was either at 0 mM or 2 mM. Complexes were incubated for 3 h at room temperature with a range of concentrations of VCA, CA, actin-VCA, GST-VCA, or a fixed concentration of polymerized actin with a titration of VCA. Complexes were incubated overnight (~12 h) with a range of concentrations of actin filaments). FlAsH-EDT_2_ was excited at λ 50 nm and emitted light was collected with a wavelength scan from 520 nm to 650 nm. Changes in donor intensity of double-labeled complexes were calculated by comparing the average fluorescence obtained from λ 530 nm to λ 532 nm (emission peak) to single-labeled complexes.

Energy transfer efficiency, ET_eff_, is defined as:

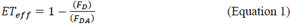

Where *F*_*D*_ is donor alone and *F_DA_* is donor with the acceptor.

We recalculated energy transfer from the data of Goley et al. (Goley, Rodenbusch et al. 2004) using Equation 2:

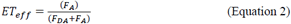

Where *F*_*A*_ is emission of the acceptor alone and *F_DA_* is the emission of the donor in the presence of the acceptor.

To transform distances (r) obtained from AV simulations to ET_effs_, ET_eff_ was defined as:

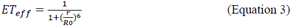

Where *r* is distance and *Ro* is the F′ster radius. R0 for FRET pair Alexa 488/594 is 54 Å (Schuler, Lipman et al. 2002)

### Total Internal Reflection Fluorescence (TIRF) Microscopy

We used a prism-style TIRF microscope to acquire fluorescence micrographs at 5s intervals with a Hamamatsu C4747-95 CCD (Orca-ER) camera controlled by MetaMorph software (Molecular Devices) on an Olympus IX-70 inverted microscope (Olympus) (Amann and Pollard 2001, Kuhn and Pollard 2005). Images were processed using ImageJ software (Schneider, Rasband et al. 2012).

Glass flow chambers (Amann and Pollard 2001) were incubated with 0.5 % Tween 20, 250 nM *N*-ethylmaleimide-inactivated skeletal muscle myosin, and twice with 5% BSA (w/v), all in TBS_HS_ buffer (50 mM Tris-HCl, 600 mM KCl, pH 75) for 3 min each. Washes for 5 min with TBS_HS_ were made after each incubation step. Chambers were equilibrated with TIRF buffer (10 mM imidazole, 50 mM KCl, 1 mM MgCl_2_, 1 mM EGTA, 50 mM DTT, 0.2 mM ATP, 0.2 μM CaCl_2_, 15 mM glucose, 20 μg/ml catalase, 100 μg/ml glucose oxidase, and 0.5 % methylcellulose (4,000 cP). 0.5 μM actin (30% labeled with Alexa 488) was polymerized in the chamber until short filaments were visible (2-5 min). Unpolymerized monomers were washed out and a mixture containing 100 nM Arp2/3 complex, 500 nM VCA, and fresh actin monomers was added during continuous imaging.

### Single molecule FRET measurements

smFRET measurements were made with an instrument built on an inverted Olympus IX-71 microscope (Olympus) using instrument settings previously described in detail (Trexler and Rhoades 2009, Metskas and Rhoades 2015). The laser power at 488 nm was set to 25 -35 μW before entry into the microscope. Fluorescence emission was collected through the objective and separated from the excitation light by a Z488RDC long-pass dichroic and 500 nm long-pass filter (Chroma). Further separation of donor and acceptor photons was achieved with HQ585LP dichroic, with an ET52550M band-pass emission filter for donor photons and a 605LP emission filter for acceptor photons. Subsequently, fluorescence emission was focused onto the aperture of an optical fiber with a 100-μm diameter (OzOptics), which was directly coupled to an avalanche photodiode (Perkin-Elmer) (Metskas and Rhoades 2015). Experiments were done in eight-well chambered cover glasses (Nunc) passivated with polylysine-conjugated polyethylene glycol which prevents proteins from sticking to chamber surfaces (Middleton and Rhoades 2010). Wells contained 75 pM labeled Arp2/3 complex with an additional 40 nM of unlabeled complex in KMET buffer and ligands described in each experiment. Under these conditions, Arp2/3 complex was stable for the duration of our measurements. Traces were collected in 1 ms time bins for at least one hour, and with only photon bursts of >30 photons per time bin included in the final histogram.

ET_eff_ values were calculated according to Equation 4

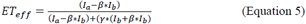

were ET_*eff*_ is energy transfer efficiency, *I*_*a*_ is the acceptor fluorescence intensity, *I*_*b*_ is the donor fluorescence intensity, *β* is a measurement of donor photon bleed-though into the acceptor channel, and ? accounts for differences in detection efficiency and quantum yield between the fluorophores. The b and g parameters are instrument-specific and were measured empirically 03B20.06, g = 1.3).ET _eff_ values were compiled into histograms which were fitted with two Gaussian distributions using MATLAB (MathWorks) scripts (Metskas and Rhoades 2015).

We estimated the upper limit for shot noise using the following formula

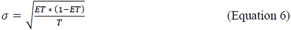

Where T is the number of photons threshold that defines a smFRET event, we fixed this value at 30 photons. ET is energy transfer obtained in our measurements.

### Alternating laser excitation (ALEX) measurements

ALEX measurements were made with an instrument based on an Olympus IX-71 inverted microscope (Olympus) (Trexler and Rhoades 2009, Metskas and Rhoades 2015). Samples were exposed to 100 μs pulses of 488 nm and 561 nm laser light controlled by acousto-optic modulators (Isomet) with 5 μs dark intervals for 60 min. Traditional ET _eff_ values and a stoichiometry (*S*) ratio were obtained using Equation 4,

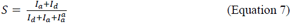

The *S* ratio characterizes labeling. Where, I_a_ is the intensity of the acceptor channel during 488 nm excitation, I_b_ is the intensity of the donor channel during 488 nm excitation, and 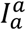 is the fluorescence intensity of the acceptor channel during 561 nm excitation. Data were analyzed with Lab-view software running an in-house script (Metskas and Rhoades 2015).

### Negative Staining

Carbon-coated copper girds (Electron Microscopy Sciences, catalog No.: CF400-CU) were glow discharged using a homemade glow discharge unit (Aebi, U. and Pollard, T.D., 1987). To absorb Arp2/3 complex to the glow discharged girds, 3.5 μL of 0.05 μM (0.011 mg/mL) Arp2/3 in KMEI buffer (50 mM KCl, 1 mM MgCl_2_, 1 mM EGTA, 10 mM imidazole, pH 7.0) or KMEI buffer with 0.2 mM ATP was incubated on the grids at room temperature for 30 s. Grids were stained with three drops of freshly prepared 0.8% (w/v) uranyl formate, and dried in air.

### Electron Microscopy

Specimens were inspected with a Tecnai F20 electron Microscope (FEI Company) operating at 200 kV. Images were recorded on a K2 Summit camera (Gatan Company) in counted mode at a nominal magnification of 29,000×, corresponding to a calibrated pixel size of 1.25 Å. The dose rate on the camera was set to ~8 counts/pixel/s, and the total exposure time was 12 s. Each image was fractionated into 20 frames (0.6 s/frame). The images were recorded at a defocus between -0.6 μm and -3.0 μm, and at a drifting speed of less than 0.2 nm/second.

### 2D Image processing

Dose-fractionated image stacks were motion-corrected and summed with MotionCorr (Li, Mooney et al. 2013). Summed images were binned by a factor of 2, resulting a pixel size of 2.5 Å. Particles were interactively picked using e2boxer in EMAN2 (Tang, Peng et al. 2007) using a box size of 120 pixels. Particle coordinates were exported from EMAN2 and imported into RELION2 (Scheres 2012) for 2D analysis. CTF estimation was performed using CTFFIND4 ((Rohou and Grigorieff 2015)), and particles were CTF corrected before reference-free 2D classification. For presentation purpose, the class averages were first sorted based on class distributions in RELION2, and then rotated to the same orientation using a cross-correlation-coefficient-based method, clipped to 100×100 pixels and masked with a radius of 40 pixels in EMAN2.

**Table 1.** Energy transfer efficiency between probes measured by smFRET

| Construct<br>Ligand | **Arp2<sub>cys</sub>-Arp3<sub>cys</sub>** | **ArpC3<sub>cys</sub>-ArpC1<sub>cys</sub>** |
| --- | --- | --- |
| None | 0.48 ± 0.01<br>(n=7) | 0.25 ± 0.02<br>(n=9) |
| + VCA | 0.61 ± 0.01<br>(n=4) | 0.28 ± 0.01<br>(n=4) |
| + Actin<sub>(m)</sub>-VCA | 0.54 ± 0.03<br>(n=3) | 0.27 ± 0.01<br>(n=6) |
| + CA | 0.60 ± 0.01<br>(n=3) | 0.28 ± 0.01<br>(n=3) |

## Acknowledgements

Research reported in this publication was supported by National Institute of General Medical Sciences of the National Institutes of Health under award number R01GM026338, an International Fellowship from the American Association of University Women, and the Yale Integrated Graduate Program in Physical and Engineering Biology. The content is solely the responsibility of the authors and does not necessarily represent the official views of the National Institutes of Health. The authors thank Shih-Chieh Ti, for providing the *S. pombe* strains used in the ensemble FRET experiments and his advice on the project.

## Competing financial interests statement (if any)

None to report.

Correspondence and requests for materials should be addressed to

